# Entraining stepping movements of Parkinson’s patients to alternating subthalamic nucleus deep brain stimulation

**DOI:** 10.1101/2020.08.16.253062

**Authors:** Petra Fischer, Shenghong He, Alexis de Roquemaurel, Harith Akram, Thomas Foltynie, Patricia Limousin, Ludvic Zrinzo, Hayriye Cagnan, Peter Brown, Huiling Tan

**Author notes:** These authors contributed equally (Petra Fischer, Shenghong He, Huiling Tan). **Corresponding authors:** Huiling Tan, Petra Fischer.

## Abstract

Patients with advanced Parkinson’s can be treated by deep brain stimulation of the subthalamic nucleus (STN). This affords a unique opportunity to record from this nucleus and stimulate it in a controlled manner. Previous work has shown that activity in the STN is modulated in a rhythmic pattern when Parkinson’s patients perform stepping movements, raising the question whether the STN is involved in the dynamic control of stepping. To answer this question, we tested whether an alternating stimulation pattern resembling the stepping-related modulation of activity in the STN could entrain patients’ stepping movements as evidence of the STN’s involvement in stepping control. Group analyses of ten Parkinson’s patients (one female) showed that alternating stimulation significantly entrained stepping rhythms. We found a remarkably consistent alignment between the stepping and stimulation cycle when the stimulation speed was close to the stepping speed in the five patients that demonstrated significant individual entrainment to the stimulation cycle. Our study provides evidence that the STN is causally involved in dynamic control of step timing, and motivates further exploration of this biomimetic stimulation pattern as a basis for the development of specific deep brain stimulation strategies to ameliorate gait impairments.

## Introduction

Some of the most challenging symptoms for patients with Parkinson’s disease are gait and balance problems as they can cause falls (Bloem, Hausdorff, Visser, & Giladi, 2004; Walton et al., 2015), loss of mobility and strongly reduce patients’ quality of life (Walton et al., 2015). Deep brain stimulation of the subthalamic nucleus (STN) is an effective treatment for tremor, rigidity and bradykinesia in Parkinson’s disease (Kleiner-Fisman et al., 2006). However, the impact of STN DBS on gait control is less consistent and can even result in deterioration of gait (Barbe et al., 2020; Collomb-Clerc & Welter, 2015). Conventional high-frequency DBS is provided continuously and is thought to attenuate beta activity (Kühn et al., 2008). Several reports describe changes in STN beta activity or its phase locking between hemispheres during gait (Arnulfo et al., 2018; Hell, Plate, Mehrkens, & Bötzel, 2018; Storzer et al., 2017), and our previous work has shown rhythmic modulation of STN activity when patients perform stepping movements (Fischer et al., 2018): Beta (20-30 Hz) activity briefly increased just after the contralateral heel strike during the stance period, resulting in alternating peaks of right and left STN activity. Auditory cueing, which also helps improve gait rhythmicity, further enhanced this alternating pattern (Fischer et al., 2018). However, whether such patterning helped organise the stepping behaviour or was secondary and afferent to it could not be discerned. Here we investigate whether STN activity is causally important in the dynamic control of stepping by assessing the entrainment of stepping by alternating high-frequency stimulation delivered to the two nuclei at a given individual’s preferred stepping speed. We also studied whether their stepping speed could be manipulated by accelerating the rhythm of alternating stimulation.

## Materials and methods

### Participants

We recorded 10 Parkinson’s patients (mean age 67 ± (STD) 7 years, disease duration 14.2 ±4 years, time since DBS implantation 3.8 ± 1.3 years, 1 female) with chronically implanted STN DBS electrodes, who had received DBS surgery 1-5 years previously at University College London Hospital (UCLH) in London (n=9) or at the Hadassah Hospital in Jerusalem, Israel (n=1). All patients were implanted with the Medtronic Activa-PC neurostimulator and the 3389 macroelectrode model to alleviate their motor symptoms, and all patients were recorded in the UK. We considered patients younger than 80 years for this study. None of the participants had cognitive impairments, which were assessed with a mini mental score examination (≥26/30 see **Table 1**). Interleaved stimulation as a DBS setting was an exclusion criterion because the streaming telemetry system Nexus-D (Medtronic, USA) that was used to control alternating stimulation cannot deliver interleaved stimulation.

**Table 1.**
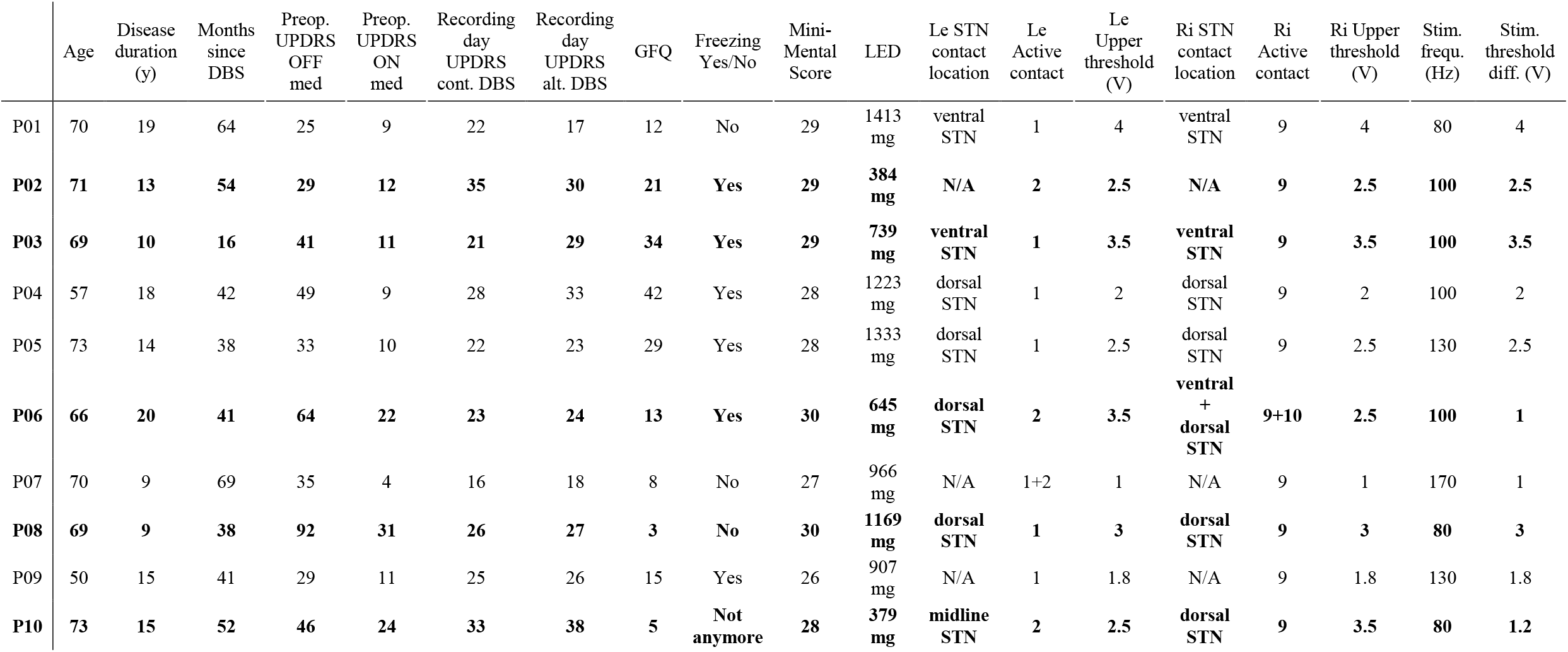
Clinical details and stimulation parameters for all patients. Patients who were significantly entrained to alternating DBS are highlighted in bold. No distinct differences between the group of responders and non-responders were apparent with respect to the stimulation intensity boundaries, location of the active contact, severity of motor symptoms or gait problems. The only criterion that stood out was the stimulation frequency, which was either 80 or 100 Hz in the group of responders. The four contacts on each electrode are labelled as 0-3 (ventral-dorsal) on the left electrode and 8-11 on the right electrode. The clinically effective stimulation intensity during standard continuous stimulation was set as *Upper threshold* (rounded to the first decimal place). *Stim threshold diff* was the difference between the upper threshold and the intensity during the periods of lower or absent stimulation during the alternating mode. This difference was the same in the two sides. All patients received stimulation with a pulse width of 60μs. GFQ = Gait and falls questionnaire (Giladi, 2000). LED = Levodopa equivalent dose.

|  | Age | Disease duration (y) | Months since DBS | Preop. UPDRS OFF med | Preop. UPDRS ON med | Recording day UPDRS cont. DBS | Recording day UPDRS alt. DBS | GFQ | Freezing Yes/No | Mini-Mental Score | LED | Le STN contact location | Le Active contact | Le Upper threshold (V) | Ri STN contact location | Ri Active contact | Ri Upper threshold (V) | Stim. frequ. (Hz) | Stim. threshold diff. (V) |
| --- | --- | --- | --- | --- | --- | --- | --- | --- | --- | --- | --- | --- | --- | --- | --- | --- | --- | --- | --- |
| P01 | 70 | 19 | 64 | 25 | 9 | 22 | 17 | 12 | No | 29 | 1413 mg | ventral STN | 1 | 4 | ventral STN | 9 | 4 | 80 | 4 |
| P02 | **71** | **13** | **54** | **29** | **12** | **35** | **30** | **21** | **Yes** | **29** | **384 mg** | **N/A** | **2** | **2.5** | **N/A** | **9** | **2.5** | **100** | **2.5** |
| P03 | **69** | **10** | **16** | **41** | **11** | **21** | **29** | **34** | **Yes** | **29** | **739 mg** | **ventral STN** | **1** | **3.5** | **ventral STN** | **9** | **3.5** | **100** | **3.5** |
| P04 | 57 | 18 | 42 | 49 | 9 | 28 | 33 | 42 | Yes | 28 | 1223 mg | dorsal STN | 1 | 2 | dorsal STN | 9 | 2 | 100 | 2 |
| P05 | 73 | 14 | 38 | 33 | 10 | 22 | 23 | 29 | Yes | 28 | 1333 mg | dorsal STN | 1 | 2.5 | dorsal STN | 9 | 2.5 | 130 | 2.5 |
| P06 | **66** | **20** | **41** | **64** | **22** | **23** | **24** | **13** | **Yes** | **30** | **645 mg** | **dorsal STN** | **2** | **3.5** | **ventral + dorsal STN** | **9+10** | **2.5** | **100** | **1** |
| P07 | 70 | 9 | 69 | 35 | 4 | 16 | 18 | 8 | No | 27 | 966 mg | N/A | 1+2 | 1 | N/A | 9 | 1 | 170 | 1 |
| P08 | **69** | **9** | **38** | **92** | **31** | **26** | **27** | **3** | **No** | **30** | **1169 mg** | **dorsal STN** | **1** | **3** | **dorsal STN** | **9** | **3** | **80** | **3** |
| P09 | 50 | 15 | 41 | 29 | 11 | 25 | 26 | 15 | Yes | 26 | 907 mg | N/A | 1 | 1.8 | N/A | 9 | 1.8 | 130 | 1.8 |
| P10 | **73** | **15** | **52** | **46** | **24** | **33** | **38** | **5** | **Not anymore** | **28** | **379 mg** | **midline STN** | **2** | **2.5** | **dorsal STN** | **9** | **3.5** | **80** | **1.2** |

The study was approved by the South Central - Oxford A Research Ethics Committee (17/SC/0416) and patients gave informed written consent before the recording.

Our main objective for this study was to find out if participants would entrain to the alternating DBS pattern and how their step timing would align to the stimulation pattern. Therefore, we did not specifically recruit patients with severe gait impairments but also included patients that experienced no gait impairments such as freezing or festination. Patients’ severity of gait impairments was assessed at the beginning of their visit with a gait and falls questionnaire (GFQ, Giladi *et al*, 2000).

### Stimulation conditions and setting the DBS parameters

All patients performed stepping in place while standing during three stimulation conditions: Conventional continuous DBS, alternating DBS at their preferred stepping speed and alternating DBS 20% faster than their preferred speed. We will refer to the latter as *fast alternating DBS* in the following sections. Some patients also performed the stepping movement when stimulation was switched off (n=5), but because time constraints allowed this only in half of all patients, this condition was not further analysed. All recordings were performed on medication to limit fatigue. Before changing DBS to the alternating pattern, patients’ preferred stepping speed was measured during ~30s free walking and during ~20s stepping in place (while DBS was on continuously) with a MATLAB script that registered the time interval between key presses performed by the experimenter at the patient’s heel strikes. Because of the highly predictable nature of the heel strike within the continuous stepping cycle, this measurement method provided a high accuracy, verified by comparing it to force plate measurements that resulted in nearly identical estimates. The key input method was chosen because it did not require any additional manual processing steps to obtain the final estimate and was thus faster. The final estimate was needed for the programming of the test conditions and was therefore needed as quickly as possible (on average, as it is, the study took 2.5 hours to complete). The key inputs were always performed by the same experimenter. The preferred duration of one full gait cycle was 1.2s in most cases (stepping in place: mean = 1.27 ± 0.22s, ranging between 1.1-1.8s, free walking: mean = 1.18 ± 0.17s, ranging between 0.94-1.4s).

There was no significant difference between the two conditions (t6 = 0.5, p = 0.664; df = 6 because the preferred speed of free walking was only measured in the final six patients). The median interstep interval from the stepping in place measurement was used to determine the duration of the stimulation cycles in the two alternating DBS conditions during stepping in place. The stimulation intensity and timing delivered by the chronically implanted pulse generator were remotely controlled by the Nexus-D device, which communicated via telemetry. The stimulation intensity was at the clinically effective voltage for two thirds of the stimulation cycle and was lowered intermittently only for one third of the full stimulation cycle (**Fig. 1A**). This rhythm was provided with an offset between the left and right STN such that the pauses occurred at opposite points within one full stimulation cycle. This 67/33% pattern was chosen because the technical limitations of Nexus-D would have not allowed a 50/50% pattern as the device requires gaps of at least 100ms to reliably send two consecutive commands (left up, right down, right up, left down, see **Fig. 1A**). We opted for 67% instead of 33% for the high-intensity stimulation period to keep the overall stimulation intensity relatively high in comparison to continuous DBS.

**Fig. 1.**
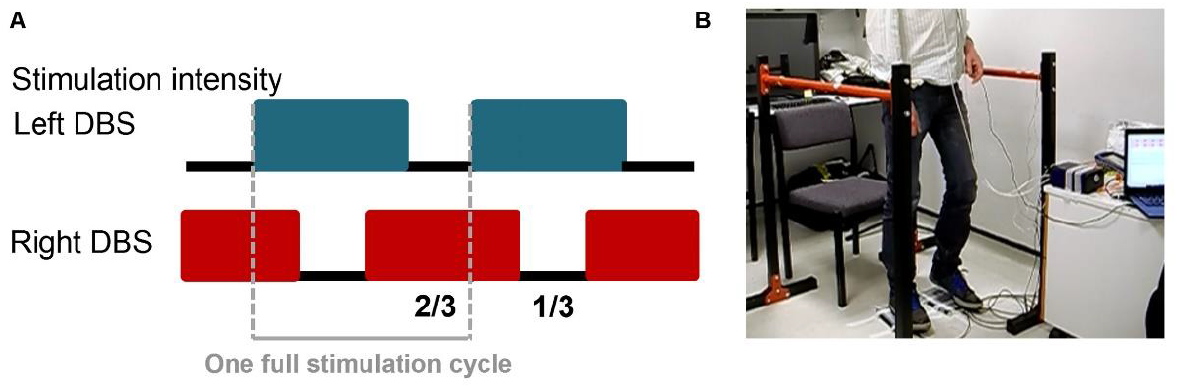
A Alternating DBS pattern. DBS was set to the clinically effective voltage for 2/3 of the stimulation cycle and reduced for 1/3 of the cycle. For the reduced period, stimulation intensity was set to 0V in eight patients and it was reduced by −1V and −1.2V relative to the clinically effective threshold in the remaining two patients. The pattern was offset between the left and right STN such that the pauses occurred at exactly opposite points of the stimulation cycle. Grey dashed lines show the start and end of one full stimulation cycle (compare with Fig. 3B). **B** Recording setup. Patients performed stepping while standing on force plates and were allowed to hold on to parallel bars positioned next to them if they felt unstable or if they felt more comfortable resting their arms on the bars.

A typical alternating stimulation cycle thus consisted of 0.8s (= 2/3 of 1.2s) of standard intensity stimulation (drawn from the clinically effective voltage during chronic continuous stimulation) and 0.4s (= 1/3 of 1.2s) of lowered intensity or no stimulation. The lower limit of alternating stimulation was determined by reducing the clinically effective voltage in steps of −0.5V and evaluating if the patient noticed a change until reaching 0V. If troublesome symptoms appeared before reaching 0V, the lower limit remained above the side effects threshold. In 8 of 10 patients the lower limit was set to 0V with patients reporting that alternating stimulation was well tolerated. In one patient (P06), reducing the lower limit by more than 1.2V resulted in reappearance of tremor and in another patient (P10) it caused headache at the forehead and slight tingling of the lips, which immediately disappeared when stimulation was switched back to the continuous mode. These two patients were the only participants with an upper stimulation threshold (based on their clinical stimulation settings) that differed between the left and the right STN (see P06 and P10 in **Table 1**). Their lower limits were set separately for the left and right STN to −1V (P06) and −1.2V (P10) below the upper thresholds, so that the patients were spared tremor and tingling. Other minor side effects in other patients were slight dizziness in one case and increased clarity, ‘as if a fog has been lifted’, in another case. Patients were informed of each change in stimulation intensity whilst the lower threshold for stimulation was sought.

Note that before using Nexus-D to switch to the alternating stimulation mode, the amplitude limits of the patient programmer option in the stimulator were adjusted with Medtronic NVision: We set the upper limit to ‘+0V’ relative to the clinical amplitude (drawn from the clinically effective voltage during chronic continuous stimulation) and the lower limit to ‘-clinical amplitude’ to ensure that the stimulation amplitude could never be increased above the clinically effective amplitude.

### Task

Patients were asked to perform stepping in place on force plates (Biometrics Ltd ForcePlates) at their comfortable speed and maintain a consistent movement throughout the recording. Two parallel bars were placed to the left and right of the force plates to allow patients to hold on to them if they wanted more stability (**Fig. 1B**). Most patients rested their arms on the bars throughout the stepping in place recordings. P02 did not use the bars, and two patients (P06 and P08) used them only intermittently as they found it less comfortable to hold on than to stand freely. The experimenter asked patients to ‘Start stepping whenever you are ready’. After about 20s they were prompted to stop and pause. For the first three patients the prompt was given verbally, and for the subsequent patients a mobile phone countdown triggered an auditory alarm after 20s to prompt the pause. The duration of the pauses was randomly varied (the shortest pause was 2.7s) and they could extend up to several minutes as patients were allowed to sit down and rest between the 20s sequences whenever they wanted. To control for any effects of fatigue that may increase with time, we chose to record the three conditions (continuous DBS, alternating DBS and fast alternating DBS) in the following order: A B C C B A, with 5-6 stepping sequences per block (except in patient P05 who completed only A B C as he was too tired to complete the full set). The order of the stimulation conditions was balanced across patients, so that A would in turn refer to continuous DBS, alternating DBS or fast alternating DBS. The stimulation was set to one mode for the whole duration of each experimental block without any pauses or resets between stepping sequences or rest intervals.

Patients were not told what stimulation condition was active. They also did not report any conscious rhythmic sensations and thus could not discern the rhythm of the alternating stimulation. The experimenter controlled the stimulation modes using custom-written software and was thus aware of the stimulation conditions but was unaware of the precise timing of the stimulation onset when prompting patients to start stepping any time again. Either before or after the stepping task, a blinded clinical research fellow performed the UPDRS-III motor examination (on medication), once during continuous DBS and once during alternating DBS. The order was randomized across patients so that continuous DBS was the first condition for half of all patients. Stepping in place provides only a proxy measure of stereotypical gait, but as part of the clinical examination a 20m free walking assessment was also performed in a corridor. For the first patients, Bluetooth communication was not yet available and one experimenter had to walk next to the patient carrying the laptop connected via USB with the Nexus-D. For the final six patients, Bluetooth communication between the laptop and Nexus-D allowed the patients to walk freely during both alternating DBS and continuous DBS. Alternating DBS was set to the individual’s preferred speed that was recorded during free walking. In these six patients, we also measured the time and number of steps needed to complete a 10m straight walk, turn and return to the starting point. Note that the step timing relative to stimulation was not recorded during free walking, and thus the strength of entrainment could not be assessed. The complete visit lasted up to 2.5 hours including extended pauses between individual assessments.

### Recordings

A TMSi Porti amplifier (2048 Hz sampling rate, TMS International, Netherlands) recorded continuous force measurements from the two force plates, which were taped to the floor, to extract the step timing. Triggers indicating the onsets of high-intensity stimulation were recorded with a light-sensitive sensor attached to the screen of the laptop that controlled stimulation timing via the Nexus-D. The screen below the sensor displayed a grey box that briefly turned black at the onset of high-intensity stimulation in the left electrode and white for the onset in the right electrode. DBS artefacts that captured if stimulation was on, and in which mode, were recorded with two bipolar electrodes attached to the back of the neck slightly below the ears. This measurement provided a simple check during the experiment that allowed us to see if the stimulation protocol was working.

### Data processing

Heel strikes were identified in Spike2 (Cambridge Electronic Design Limited) based on the force measurements by setting a threshold for each patient to capture approximately the midpoint of each force increase (**Fig. 2**). The force measurement increased whenever weight was transferred onto a force plate. Note that the foot touched the force plate already slightly earlier, about 100ms before the heel strike event, however, considerable weight was only transferred on the leg by the time of the event. We used the same threshold for identifying when the leg was lifted, which was captured by a force decrease. Note here again that the foot was fully lifted off the plate only slightly after the event, however, the process of lifting the leg up was initiated already before then.

**Fig. 2.**
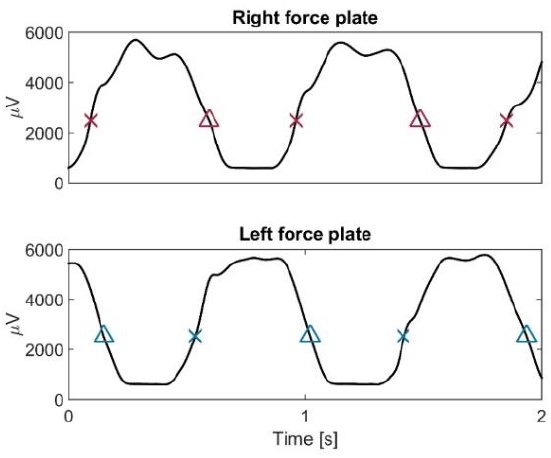
Force measurements and step cycle events. x = heel strikes. The force increased during heel strikes. Δ = when the foot was raised from the force plate the force decreased.

To avoid biasing the entrainment results by sequences that were several seconds longer than other sequences, which occurred occasionally when verbal prompts were used to prompt stopping, steps at the beginning and end of the longer sequences were removed, such that the remaining number of steps did not exceed the median number of steps of all the sequences.

Freezing episodes were very rare and were excluded from the analyses. They occurred in two patients (P03, P04) towards the end of the recording session without any apparent difference between conditions.

### Statistical analysis

All analyses were performed with MATLAB (v. 2016a, The MathWorks Inc., Natick, Massachusetts). Here we define entrainment as significant alignment of the timing of steps to the rhythm of the alternating stimulation pattern. This alignment was evaluated with a Rayleigh-test (using the MATLAB toolbox CircStat; Berens, 2009) for each individual patient and with a permutation procedure at the group level that considers each individual’s average timing and entrainment strength.

Whenever a heel strike occurred (tests are only reported for the left heel strikes, because p-values were highly similar for the right heel strike), the coincident phase of the rhythmic alternating DBS pattern was extracted. The uniformity of this resulting phase distribution was then assessed with a Rayleigh-test to test if individual patients showed significant entrainment. An additional permutation procedure was used to compute a group statistic across all ten recorded patients. For the group statistic, the vector length was calculated first for each patient according to the formula 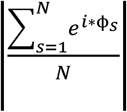, where ϕ_*s*_ is the phase of alternating DBS at each left heel strike and *N* the number of all heel strikes. The grey dashed lines in Fig. 1A show the start and end of one full stimulation cycle, and the x-axis in Fig. 3B shows the phase of one alternating stimulation cycle. Note that whenever we show arrows representing phases, they always refer to the phase of alternating stimulation at the time of the patients’ heel strikes and not to the phase of their stepping cycle, which was another cyclic measurement. The circular mean of these phases was then computed to obtain the average ‘preferred’ phase for each patient. This resulted in ten vectors (one for each patient) with their direction representing the average preferred phase, and their length representing the strength of entrainment (blue vectors in **Fig. 3A**). Next, they were transformed into Cartesian coordinates and the average of the ten vectors (black vector in **Fig. 3A**) was computed. The length of this average vector was obtained using Pythagoras’ theorem and was our group statistic of interest. It takes into account both the strength of entrainment and the consistency of the preferred phases across patients. If all patients would have shown strong entrainment, but with different preferred phases, the length of the group average vector would be close to zero. Only if the vectors representing individual patients pointed into a similar direction, the group average vector would be significantly larger than the one obtained from our permutation data.

**Fig. 3.**
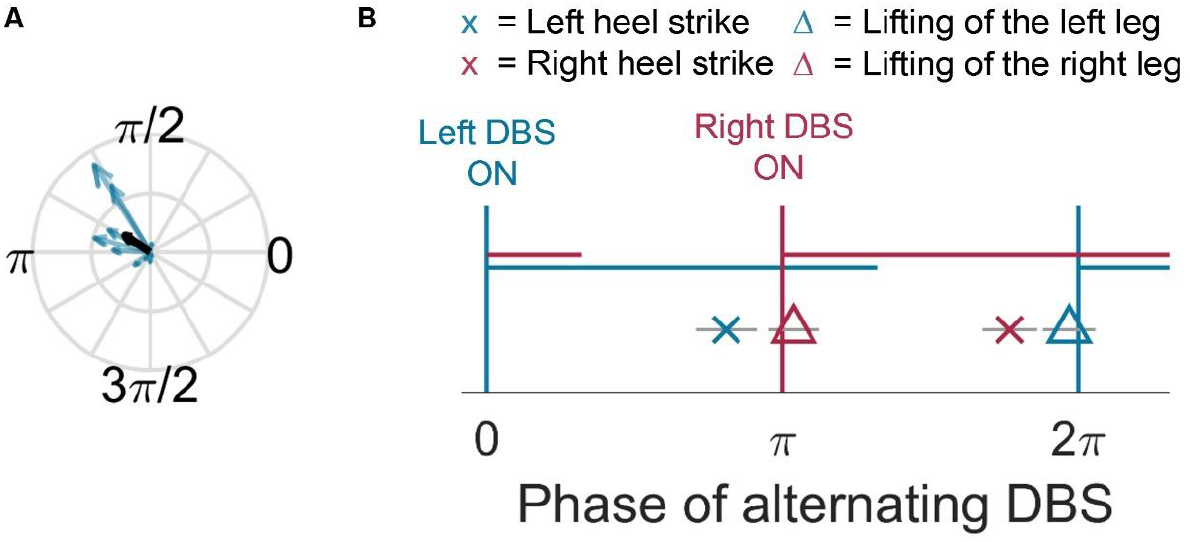
Entrainment at the group level. **A** Blue vectors show the average phase of alternating DBS at all left heel strikes and the strength of entrainment for individual patients (n=10). Long arrows show strong entrainment. The group average vector (black arrow) shows the average of the blue vectors. The length of this vector was significantly larger than in the surrogate data, demonstrating consistent alignment of stepping to the alternating DBS pattern across the group. **B** Group-averaged timing of key events of the gait cycle (x and Δ) relative to the stimulation pattern. The blue and red horizontal lines indicate high-intensity stimulation of the left and right STN, respectively. The left heel strike (blue x) was made just before contralateral stimulation (right STN DBS shown in red) increased. Grey horizontal bars indicate the standard error of the mean phases across the patients.

We computed a permutation distribution of 1000 surrogate vector lengths by shifting, separately for each patient, each of their 20s long stepping sequences in time by a random offset drawn from a uniform distribution ranging between −1.5s and +1.5s. This way the rhythmic structure within the 20s stepping sequences remained intact and only their relative alignment to the stimulation pattern was randomly shifted. Once all sequences were randomly shifted, we computed the surrogate vector length and preferred phase for each patient as described above for the unpermuted data. The resulting ten surrogate vectors were again averaged in the Cartesian coordinate system to compute the average length as described above. After repeating this 1000 times, we obtained a p-value by counting how many of the surrogate group vector lengths (L_*p*_) were larger or equal to the original group vector length (L_*orig*_) and dividing this number by the number of permutations (*N_p_*). The number 1 is added to both the nominator and the denominator to avoid p-values of 0 and be consistent with the exact p-value, which must be at least 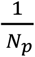 (see section 4.2 from Ernst, 2004):

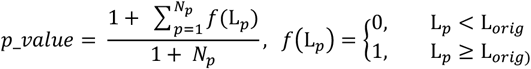

As we expected entrainment to be strongest when the stimulation speed matches the patient’s stepping speed as closely as possible, the group statistic was based on the data from the alternating DBS condition that matched the patient’s stepping speed most closely. All patients that showed significant entrainment indeed did so in the condition that was closest to their stepping speed. The stepping pace of several patients (P03-P08) was considerably faster during the recording than in the brief initial assessment, hence in those, the fast alternating DBS condition matched their performed stepping rhythm more closely.

Pairwise comparisons of the step intervals between the two alternating DBS conditions and of the change in variability between speed-matched alternating DBS and continuous DBS were performed using two-tailed t-tests or Wilcoxon signed-rank tests (with an alpha-level of 0.05) if the normality assumption (assessed by Lilliefors tests) was violated. To get a robust estimate for each patient and condition, first the median of all step intervals within each 20s stepping sequence was computed, and then again the median over all sequences was computed. To investigate the step timing variability, we computed the coefficient of variation of the step intervals (STD / mean * 100) as well as the standard deviation of the difference between two consecutive step intervals for each sequence. The median over all sequences was again computed to get a robust estimate.

To test in each patient individually if the step timing variability was significantly modulated by alternating DBS, we computed two-samples t-tests or rank-sum tests (if the normality or variance homogeneity assumption was violated) between the step timing variability estimates of the stepping sequences that were recorded in each DBS condition.

### Localization of the active electrode contacts

Each DBS lead has four contacts of which only one or two are activated during stimulation. The location of the active contacts was assessed in Brainlab (Brainlab AG, Germany) by a neurosurgeon and a neurologist who manually drew the lead on the post-operative T1 MR images centered on the DBS electrode artefact. The position of the contacts within the STN was then assessed visually in the patients’ pre-operative artefact-free T2 images. We did not have access to imaging data for P7 who received the surgery in Israel, and the quality of the imaging data was insufficient in two patients, so in these cases no accurate estimate of the contact position could be obtained.

### Data availability

The data that support the findings of this study and custom code used for analyses are available from the corresponding author upon request.

## Results

### Entrainment to DBS which alternates with a frequency matching that of stepping

Ten patients with Parkinson’s disease started sequences of 20s stepping in place while alternating DBS was already ongoing. Testing for significant entrainment of their steps to the stimulation pattern thus quantified to which extent patients aligned their stepping rhythm in each sequence to the ongoing stimulation pattern despite not being consciously aware of the precise pattern. An example of the recorded force plate measurements is shown in **Fig. 2**. **Fig. 3A** shows significant entrainment of the stepping movement to altDBS at the group level compared to surrogate data (p=0.002). The fact that all long vectors point into the same corner highlights that the preferred phase was remarkably consistent across patients. We also confirmed this finding using a simple Rayleigh test, comparing the preferred phases across patients irrespective of the strength of their entrainment, as this cannot be taken into account by a conventional Rayleigh-test. This demonstrated again significant clustering of three of the four stepping events (left heel strike p = 0.109, right heel strike: p = 0.033, left leg raised: p = 0.020; right leg raised: p = 0.015).

On an individual level, half of the ten recorded patients showed significant entrainment in the speed-matched stimulation condition (**Table 2**). **Fig. 4A** shows two examples of patients that were significantly entrained and **Fig. 4B** shows one example of a patient that was not entrained. The two plots to the left show the stimulation phases coinciding with the left and right heel strikes. The plots to the right with fewer arrows show the preferred phase and strength of entrainment for each of the separate sequences of 20s stepping that patients performed. The arrows are clustered again around the preferred phase in the patients that were entrained to the stimulation pattern, which was not the case in **Fig. 4B**. **Table 1** shows the stimulation parameters and location of the electrode contact used for stimulation. The location of the active contacts varied across patients such that some were located in the ventral, some in the dorsal STN, but no criteria emerged that would distinguish between the groups of responders. The only parameter that may be associated with entrainment may be the stimulation frequency, as in the group of responders it was either 80 Hz or 100 Hz, but never 130 Hz, which is the conventional frequency for STN DBS (Moro et al., 2002). However, two non-responders also had a stimulation frequency of 80 and 100 Hz.

**Fig. 4.**
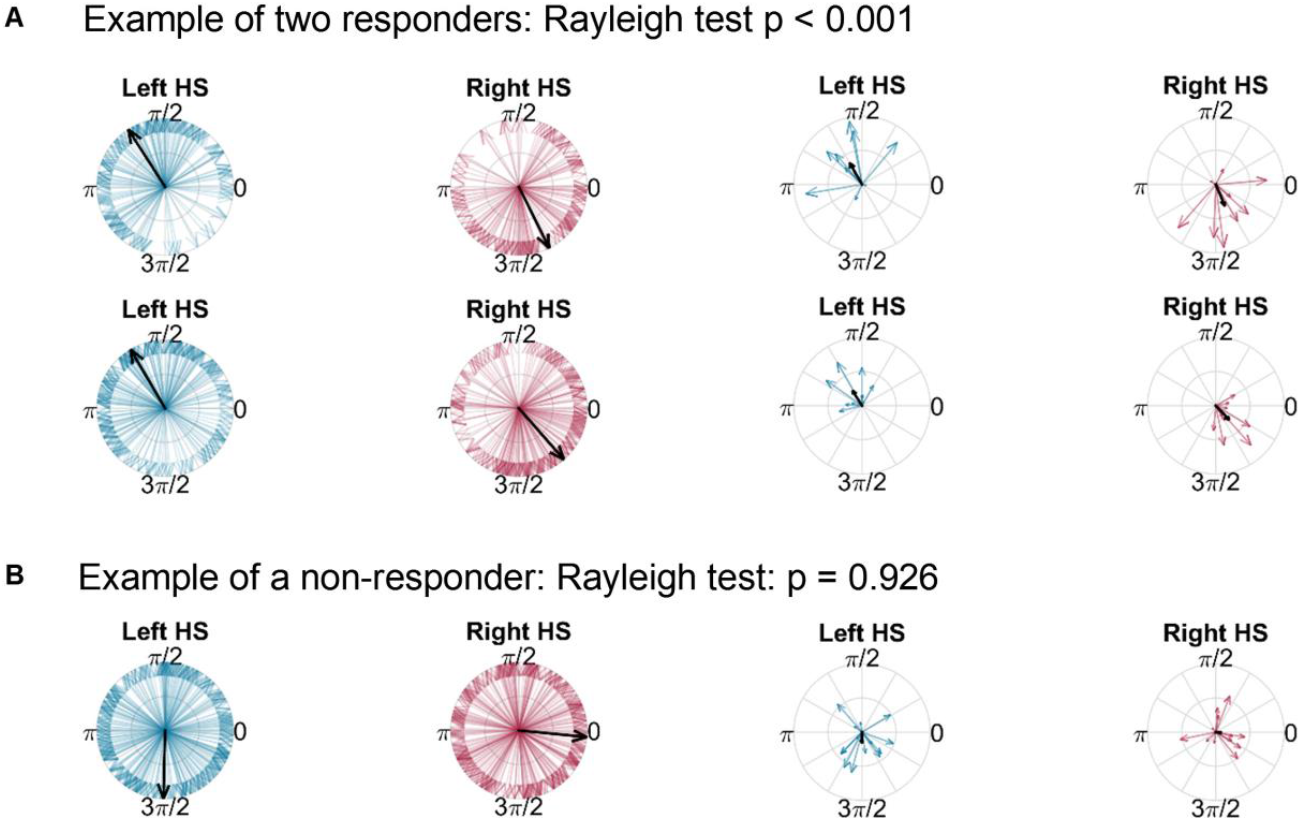
A Example data of two responders (P02 and P03). Blue and red vectors show the phases of the alternating stimulation pattern at the time of the left and right heel strikes, respectively. The heel strikes were clustered around one point of the stimulation cycle (between ∏/2 and ∏ for the left heel strike). The black vectors show the average preferred phase (scaled to unit length on the left two plots to enable a better visual comparison of the similarity between the two patients). The two plots to the right show the preferred phase and strength of entrainment (indicated by the length of the black vector) for each of the separate sequences of 20s stepping. Here the vectors also point relatively consistently to the same quarter. **B** No consistent clustering was present in non-responders (P04).

**Table 2.**
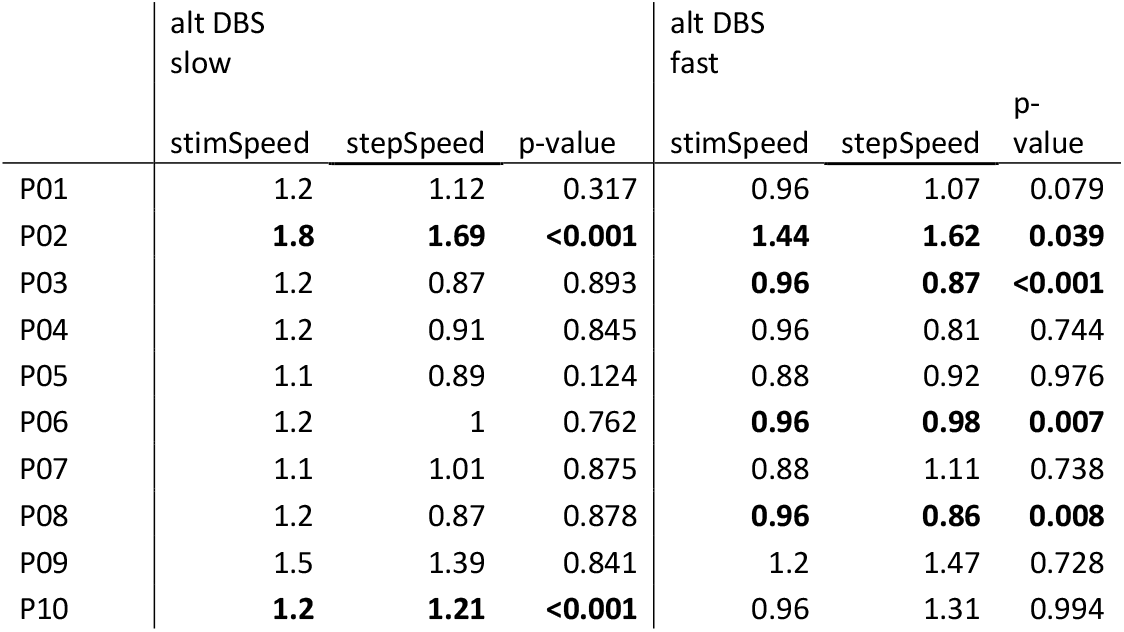
Stimulation speed, stepping speed and p-values testing for significant entrainment in the two alternating DBS conditions. The p-values in bold highlight the patients that were significantly entrained to the alternating DBS pattern (assessed with Rayleigh-tests). Significant entrainment always occurred in the condition in which the stepping speed was closer to the stimulation speed (sometimes this was in the altDBS fast condition as some patients performed the task faster than in the initial speed recording). Only P02 was also entrained to alternating DBS in the other condition. P05 and P07 reported that when stimulation was switched off outside of this study, they did not notice an immediate deterioration of symptoms, suggesting that DBS only had weak positive effects. These two patients were not entrained to alternating

|  | alt DBS<br>slow |  |  | alt DBS<br>fast |  |  |
| --- | --- | --- | --- | --- | --- | --- |
|  | stimSpeed | stepSpeed | p-value | stimSpeed | stepSpeed | p-value |
| P01 | 1.2 | 1.12 | 0.317 | 0.96 | 1.07 | 0.079 |
| P02 | **1.8** | **1.69** | **<0.001** | **1.44** | **1.62** | **0.039** |
| P03 | 1.2 | 0.87 | 0.893 | **0.96** | **0.87** | **<0.001** |
| P04 | 1.2 | 0.91 | 0.845 | 0.96 | 0.81 | 0.744 |
| P05 | 1.1 | 0.89 | 0.124 | 0.88 | 0.92 | 0.976 |
| P06 | 1.2 | 1 | 0.762 | **0.96** | **0.98** | **0.007** |
| P07 | 1.1 | 1.01 | 0.875 | 0.88 | 1.11 | 0.738 |
| P08 | 1.2 | 0.87 | 0.878 | **0.96** | **0.86** | **0.008** |
| P09 | 1.5 | 1.39 | 0.841 | 1.2 | 1.47 | 0.728 |
| P10 | **1.2** | **1.21** | **<0.001** | 0.96 | 1.31 | 0.994 |

### Faster alternating DBS did not systematically accelerate patients’ stepping rhythm

We also tested if patients’ stepping rhythms were faster in the fast altDBS condition compared to the slower altDBS condition. We performed this comparison across all patients to test if speeding up the stimulation pattern would generally accelerate the stepping rhythm, irrespective of which condition matched their speed more closely. **Fig. 5** shows that the stepping intervals were not systematically shortened (left plot, altDBS = 0.55 ± 0.13s, fast altDBS = 0.55 ± 0.14s, t(9) = −0.3, p = 0.806). We also compared the change in interval duration relative to the baseline condition of continuous DBS, which again showed that the fast DBS condition resulted in speed changes in either direction (**Fig. 5**, right plot).

**Fig. 5.**
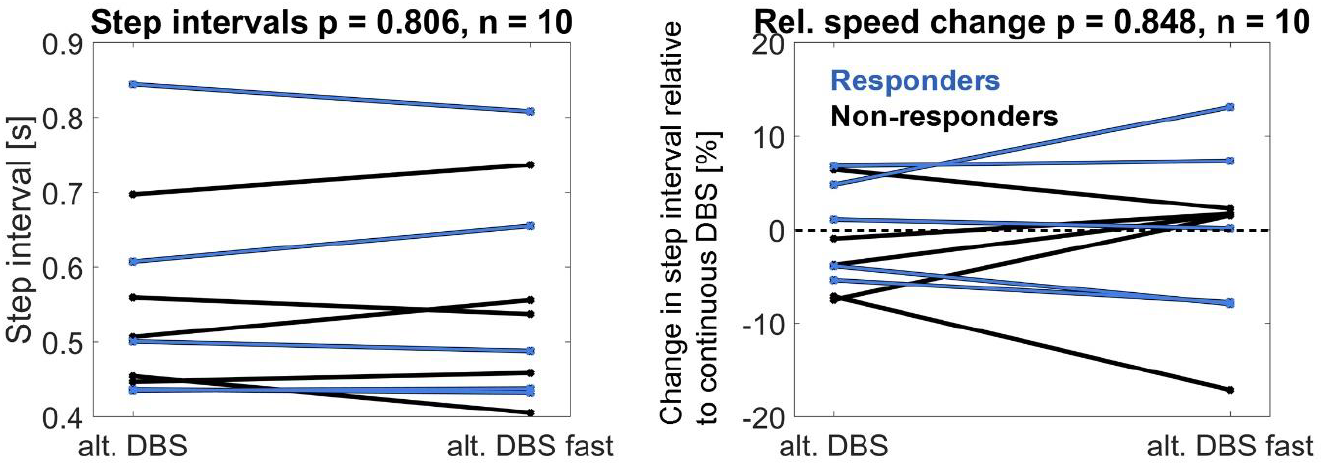
Difference in step intervals between the alternating DBS and the fast alternating DBS condition. When the alternating DBS rhythm was 20% faster, the stepping intervals were not systematically accelerated.

We also looked for order effects and found no evidence of these on stepping speed or the strength of entrainment in the speed-matched and fast-alternating conditions. In three responders (P06, P08 and P10) the two alternating DBS conditions were separated by the continuous DBS condition, showing that the strength of entrainment was not dependent on potentiation effects of prolonged alternating stimulation.

### Step timing variability during alternating DBS

First, we compared if the step timing variability changed in the alternating speed-matched DBS condition compared to continuous DBS. The variability metrics were computed within stepping sequences that included on average 40 ±5 steps. No significant differences were found across the ten patients in the coefficient of variation (CV) of the step intervals (contDBS = 8.3 ±3.4%, speed-matched altDBS = 9.3 ±3.2%, t(9) = −0.8, p = 0.450) or in the STD of the differences between consecutive step intervals (contDBS = 0.07 ±0.03, speed-matched altDBS = 0.07 ±0.03, t(9) = −0.4, p = 0.674).

Next we restricted the analysis to the group of responders, and found that the CV of the step intervals in the speed-matched alternating DBS condition was increased compared to continuous DBS (contDBS = 8.2 ±3.0%, speed-matched altDBS = 10.9 ±3.9%, t(4) = −2.9, p = 0.045). This is consistent with a failure of the step cycle to continuously entrain to the alternating stimulation rhythm, leading to increased phase slips as stepping falls in and out of register with the stimulation rhythm. When testing individually in each patient how the step timing variability changed between the stepping sequences recorded in the contDBS and speed-matched altDBS conditions, two of the five patients showed significantly increased variability during alternating DBS (rank-sum test between the respective stepping sequences: P03 p = 0.040, P08: p = 0.004).

In the group of the five responders, we also compared if their step timing variability differed between the speed-matched and mismatched altDBS condition. We found no significant difference across the group (speed-matched altDBS = 10.9 ±3.9%, mismatched altDBS = 9.9 +2.9%, t(4) = 2.1, p = 0.101), but in the within-patients tests, one of the responders (P10) had a significantly higher step timing variability when stimulated with mismatched altDBS compared to speed-matched altDBS (two-samples t-test: t(21) = −2.8, p = 0.010).

### Clinical assessments

The blinded UPDRS-III assessment showed no significant differences between continuous DBS (25.1 ± (STD) 5.7) and alternating DBS at the preferred walking speed (26.5 ± 6.45, Wilcoxon signed-rank test (n=10), p = 0.254). The UPDRS items 27-31 reflecting balance and gait also were very similar (in seven of the ten recorded patients the scores were identical between conditions, and p-values of the signed-rank tests were 1.0; item 27 mean: contDBS = 0.8 ±0.6, altDBS = 0.9 ±0.9; item 28: contDBS = 0.8 ±0.6, altDBS = 0.9 ±0.9; item 29: contDBS = 1.2 ±0.4, altDBS = 1.2 ±0.4; item 30: contDBS = 1.0 ±0.7, altDBS = 1.1 ±0.9; item 31: contDBS = 1.4 ±0.5, altDBS = 1.5 ±0.7). In the six patients that performed a timed 20m walking assessment (walk 10m straight, turn and return back to the starting point) the time needed and numbers of steps did not differ significantly between stimulation conditions (continuous DBS: 19.8s ± 5.2s and 35 ±8 steps, alternating DBS: 19.8s ± 4.5s and 35 ±6 steps).

## Discussion

We found that alternating DBS – intermittently lowering and increasing stimulation intensity with an offset between the right and left STN to produce an alternating stimulation pattern – can significantly manipulate the step timing of Parkinson’s patients. The preferred timing of the steps relative to the stimulation pattern was highly consistent across the patients that significantly entrained to alternating DBS, providing evidence that the STN is mechanistically involved in organising stepping. This is consistent with the alternating pattern of beta activity previously reported in the STN during stepping movements (Fischer et al., 2018), although, by themselves, correlational observations so far could not distinguish between the mechanistic or secondary (afferent) involvement of STN activity (Fischer et al., 2018; Georgiades et al., 2019; Singh et al., 2013).

Our findings also suggest that entrainment only occurs when the stimulation speed closely matches the participants’ stepping speed. The faster alternating DBS condition, which was accelerated by 20%, failed to accelerate patients’ stepping speed. Amongst responders, alternating DBS could increase patients’ step timing variability. Step timing variability would not change if the stepping and stimulation rhythms were aligned only by coincidence. The increase in variability suggests that entrainment was relatively weak and that stimulation can act like an attractor, pulling the intrinsic rhythm in to register, but only intermittently, punctuated by phase slips. How frequently phase slips occur likely depends on how well the alternating stimulation rhythm matches that of natural stepping.

We would like to acknowledge that stepping in place performance does not necessarily reflect how alternating DBS would affect gait variability during free walking. Despite the instruction to maintain a comfortable stepping movement as consistently as possible, some patients showed considerable variability in how high they lifted their feet across the recording session and even within individual stepping sequences, which may have affected their step intervals.

As we had no recordings of leg kinematics, this could not be quantified or analysed further. We decided to use stepping in place on force plates for the entrainment assessment because it is safer than free walking, could be performed in a relatively small space and provided a simple measure of step timing, which was our main focus in this study. Moreover, the speed of stepping in place appears to match the speed of real walking reasonably well, at least in healthy participants (Garcia, Nelson, Ling, & Van Olden, 2001).

Our study was not optimized for testing potential therapeutic benefits of alternating DBS, but we have now attained a first template for the preferred alignment between alternating DBS and the stepping cycle based on the five responders. This template can be used to inform future studies, in which the stimulation pattern could be aligned to the stepping rhythm as the patient starts walking with the help of external cues or by tracking the stepping rhythm (Tan et al., 2018).

We chose to stimulate at a high intensity for two thirds of the gait cycle and reduce stimulation for one third of the gait cycle, partially because the device used to communicate with the implanted impulse generator did not allow a 50-50% stimulation pattern. Based on our findings, we cannot infer the preferred alignment for other stimulation patterns or if the strength of entrainment would differ.

The consistent entrainment patterns among the responders cannot be explained by an awareness of the stimulation condition because none of the patients reported any rhythmic stimulation-induced sensations when asked if anything felt different. Five of our ten patients did not get entrained to alternating DBS. Two of these patients reported that switching DBS off outside of this study did not result in immediately noticeable deterioration of symptoms, and are thus atypical in their response to DBS, but were still included in the analyses. For the remaining three patients it is less clear why their stepping was not entrained. As we did not assess how quickly motor symptoms deteriorated OFF DBS and recovered after switching it back on, we could not investigate if rapid responses to changes in DBS were linked to responsiveness to alternating DBS. The stimulation speed for the non-responders was matched similarly well to their stepping speed as in the group of responders, and the severity of gait impairments was similarly variable. The presence of freezing also did not seem to play a role in this comparatively small sample. Also the location of the active DBS contacts did not appear to be critical, considering that in some responders the active contacts were located in the dorsal while in others they were in the ventral part of the STN. The only criterion that stood out was that the patients in the responding group had a stimulation frequency of either 80 or 100 Hz, slightly lower than the conventional stimulation frequency of 130 Hz for STN DBS (Moro et al., 2002). This is interesting considering that several studies suggest that lowering the frequency can be beneficial for improving gait problems in some patients (di Biase & Fasano, 2016; Di Giulio et al., 2019; Xie et al., 2018). The question whether the stimulation frequency plays a critical role in enabling entrainment to alternating DBS should be tested in future studies.

At present we can only speculate about the mechanisms underlying the observed entrainment. Patients tended to perform the most effortful part of the gait cycle – lifting a foot off the ground – after the contralateral STN had been stimulated at the clinically effective threshold for several hundred milliseconds, which is in line with the known movement-facilitatory effects of DBS. High-intensity stimulation also coincided with the time of the beta rebound, which peaks after the contralateral heel strike according to our previous study (Fischer et al., 2018). Because STN DBS can counteract excessive beta synchrony (Eusebio & Brown, 2009; Tinkhauser et al., 2017), stimulating with a high intensity after the contralateral heel strike could potentially prevent beta synchronization going overboard in the stance period. Excessive beta synchrony has recently been related to freezing episodes (Georgiades et al., 2019; Storzer et al., 2017) and to the vulnerability to such episodes (Chen et al., 2019), hence stimulating more strongly at points where beta synchronization is more likely may be a more effective stimulation strategy for preventing freezing than continuous DBS.

A recent study also found that non-invasive transcranial alternating current stimulation (tACS) over the cerebellum can entrain the walking rhythm of healthy participants (Koganemaru et al., 2019). The STN projects to the cerebellum via the pontine nuclei, thus alternating STN DBS could potentially entrain the gait rhythm via this route (Bostan, Dum, & Strick, 2010). The pedunculopontine nucleus (PPN), part of the mesencephalic locomotor region, also is reciprocally connected with the STN, and might provide another pathway by which STN DBS modulates stepping (Jenkinson et al., 2009; Morita et al., 2014; Thevathasan et al., 2018). Finally, the STN also communicates with the mesencephalic locomotor region through the substantia nigra pars reticulata (Hamani, Saint-Cyr, Fraser, Kaplitt, & Lozano, 2004). The latter structure may be preferentially sensitive to lower stimulation frequencies (Weiss, Milosevic, & Gharabaghi, 2019), and it is interesting to highlight again that lower stimulation frequencies tended to be associated with successful entrainment to alternating stimulation in the present study.

In summary, this study provides evidence that the STN is causally important in the dynamic control of the stepping cycle and provides a novel means of modulating this control through alternating STN DBS in patients with Parkinson’s disease. This stimulation mode can entrain stepping and parallels the alternating pattern of beta activity recorded in the STN during gait. It remains to be seen whether such a potentially biomimetic stimulation pattern can provide the basis for a novel treatment strategy for patients with debilitating gait disturbances. Our results suggest that it will be key to match the stimulation pattern closely to the patients’ preferred walking speed if this is to be reinforced through entrainment.

## Abbreviations

DBS: Deep brain stimulation
altDBS: Alternating deep brain stimulation
contDBS: Continuous deep brain stimulation
STN: Subthalamic nucleus
UPDRS: Unified Parkinson’s Disease Rating Scale

## Acknowledgements

The authors would like to thank all the patients who have kindly participated in this study, Medtronic for supplying the Nexus-D device and Professor Timothy Denison on advising us on wireless communication protocols.

